# Dynamics and nanoscale organization of the postsynaptic endocytic zone at excitatory synapses

**DOI:** 10.1101/2021.02.18.431766

**Authors:** Lisa A.E. Catsburg, Manon Westra, Annemarie M. L. van Schaik, Harold D. MacGillavry

## Abstract

At postsynaptic sites of neurons, a prominent clathrin-coated structure, the endocytic zone (EZ), controls the trafficking of glutamate receptors and is essential for synaptic plasticity. Despite its importance, little is known about how this clathrin structure is organized to mediate endocytosis. We used live-cell and super-resolution microscopy techniques to reveal the dynamic organization of this poorly understood clathrin structure. We found that a subset of endocytic proteins only transiently appeared at postsynaptic sites. In contrast, other proteins, including Eps15, intersectin1L, and β2-adaptin, were persistently enriched and partitioned at the edge of the EZ. We found that uncoupling the EZ from the synapse led to the loss of most of these components, while disrupting the actin cytoskeleton or AP2-membrane interactions did not alter EZ positioning. We conclude that the EZ is a stable, highly organized molecular platform where components are differentially recruited and positioned to orchestrate the endocytosis of synaptic receptors.

## INTRODUCTION

Clathrin-mediated endocytosis is the principal mechanism for the internalization of membrane components, and is essential for cellular homeostasis, intercellular signaling and nutrient uptake in mammalian cells (Kaksonen and Roux, 2018; McMahon and Boucrot, 2011; Mettlen et al., 2018). This process involves the tightly-controlled initiation and maturation of clathrin-coated pits that is mediated by the sequential recruitment of clathrin, cargo and endocytic adaptor proteins (Cocucci et al., 2012; Taylor et al., 2011). Apart from these well-characterized, small (∼100 nm) and short-lived (<120 sec) clathrin coats, numerous electron and (live-cell) light microscopy studies have revealed that clathrin can assemble into a remarkably large variety of membrane-attached structures (Grove et al., 2014; Heuser, 1980; Leyton-Puig et al., 2017; Saffarian et al., 2009; Sanan and Anderson, 1991). In fact, the lifetime, size and morphology of clathrin assemblies at the membrane diverge enormously between cell types and even within cells. Clathrin structures varying from 100 nm up to 1 µm with lifetimes ranging from seconds to tens of minutes have been reported. The origins and functional relevance of this striking heterogeneity remain to be elucidated.

This heterogeneity is particularly evident in neurons that contain a divergent population of clathrin structures distributed over their immense and complex plasma membrane. At postsynaptic sites, a clathrin-coated structure referred to as the endocytic zone (EZ) is stably associated with the postsynaptic density (PSD) (Blanpied et al., 2002; Lu et al., 2007), via a Shank-Homer1c-Dynamin3 interaction (Lu et al., 2007; Rosendale et al., 2017). Disrupting the PSD-EZ interaction severely affects glutamate receptor levels at synapses. Particularly, the ionotropic AMPA-type glutamate receptors (Petrini et al., 2009; Rosendale et al., 2017) and metabotropic glutamate receptors (Scheefhals et al., 2019) have been found to undergo trafficking mediated by the EZ, while transferrin receptors are not preferentially internalized near the synapse (Rosendale et al., 2017). It has been proposed that once internalized at the EZ, glutamate receptors enter the local recycling mechanism, that retains receptors in intracellular pools that can recycle back to the synaptic membrane in an activity-dependent manner (Park et al., 2006). Indeed, the local recycling of receptors via the EZ is essential for synaptic plasticity as uncoupling the EZ from the PSD depletes synaptic AMPA receptors and aborts activity-induced trafficking of receptors to the synaptic membrane during long-term potentiation (Lu et al., 2007; Petrini et al., 2009). Importantly, disruptions in EZ structure and function have been associated with the development of neuronal disorders such as autism spectrum disorder and Parkinson’s disease (Cortese et al., 2016; Scheefhals et al., 2019).

Despite the clear functional importance of the EZ for synaptic transmission and plasticity in neurons, the molecular organization and how this organization contributes to its function is poorly understood. In electron microscopy studies, clathrin-coated structures have been observed within dendritic spines (Petralia et al., 2003; Tao-Cheng et al., 2011) at an approximate distance of 100-600 nm from the PSD, coinciding with an enrichment of adaptor proteins such as dynamin2 and AP2 (Rácz et al., 2004). However, fundamental information on the spatial distribution and dynamics of endocytic proteins relative to the EZ and how these proteins contribute to EZ organization is missing. Here, we resolved the spatial and temporal organization of clathrin-coated structures in dendrites and spines using live-cell imaging and super-resolution microscopy. We found that the postsynaptic EZ contains a unique and stable assembly of endocytic proteins, that is highly organized at the nanoscale level. Based on these findings, we propose that the EZ is a highly distinct clathrin-coated structure that operates as a preassembled platform for endocytosis of synaptic components to sustain efficient synaptic transmission and plasticity.

## RESULTS

### Heterogenous morphology of clathrin-coated structures in dendrites

To visualize clathrin-coated structures in mature cultured hippocampal neurons (DIV16-21), GFP-clathrin light-chain-A (GFP-CLCa) was co-transfected with Homer1c-mCherry as a marker of excitatory synapses. We found a large variety of clathrin-coated structures distributed throughout the entire neuron (Figure 1A). In dendrites, a high density of clathrin structures was found in the shaft and the majority of dendritic spines contained a distinct EZ, defined as a clathrin puncta closely associated with the PSD (75 ± 5%), consistent with previous observations (Blanpied et al., 2002; Lu et al., 2007; Scheefhals et al., 2019). Importantly, labelling endogenous CLCa using a CRISPR/Cas9-based approach (Willems et al., 2020) resulted in comparable distribution of clathrin structures (Supplement Figure 1A-C). To resolve clathrin-coated structures in dendrites at high spatial resolution, we used stimulated emission depletion (STED) microscopy, allowing quantitative analyses of clathrin structure morphology (Figure 1B). Notably, STED resolved individual structures at much higher resolution than confocal, often resolving distinct substructures within clathrin patches that appeared homogenous in confocal microscopy (Figure 1C). PSDs associated with more than one clathrin structure were also observed (Figure 1B, C). We found that 59 ± 5% of the PSDs were associated with one clathrin structure, while 14 ± 2% and 5.4 ± 0.8% were associated with two or three clathrin structures, respectively (Figure 1C).

**Figure 1.**
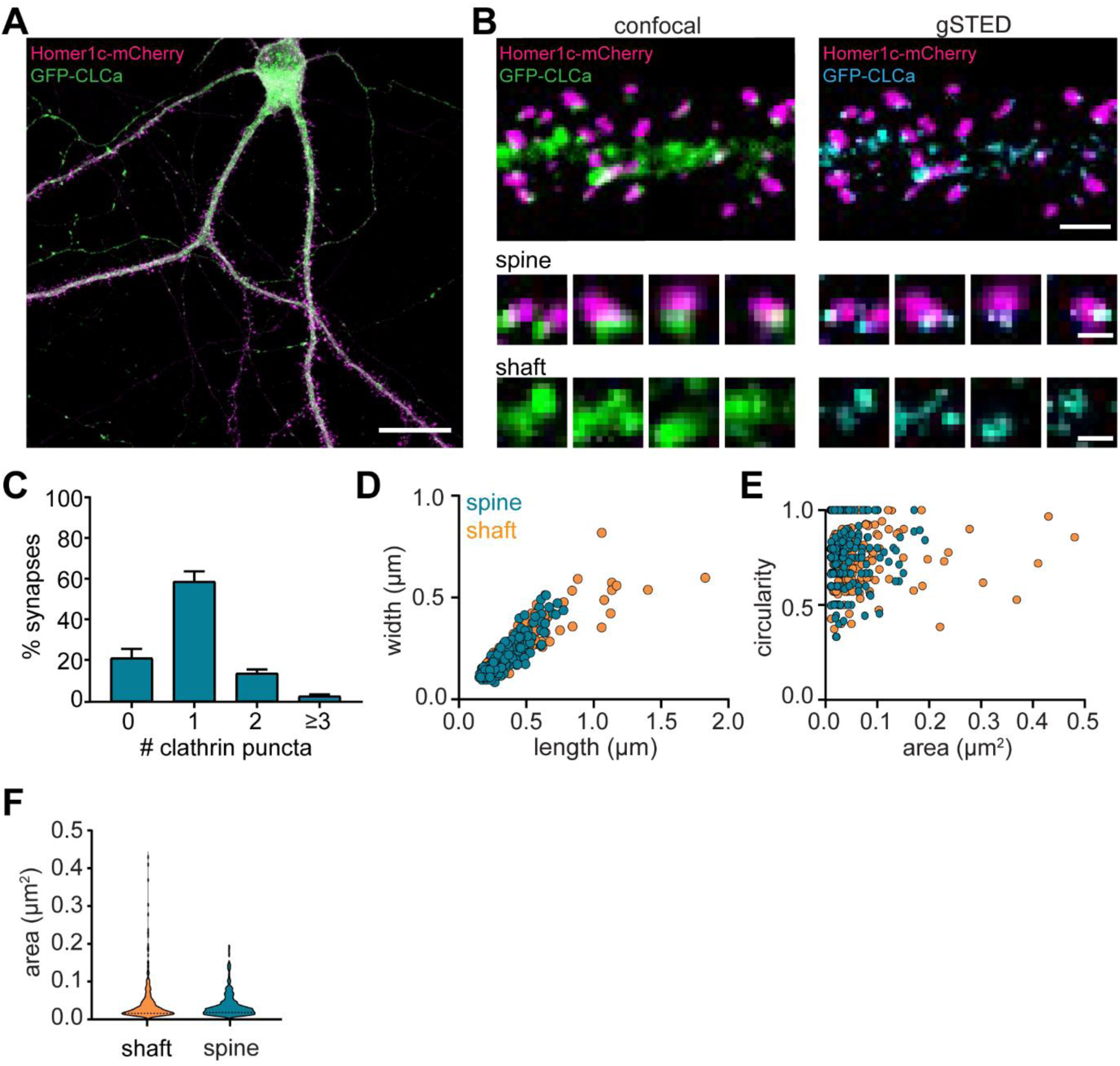
Heterogenous morphology of clathrin-coated structures in dendrites. (A) Example image of neuron expressing Homer1c-mCherry and GFP-CLCa. Scale bar: 20 µm. (B) Comparison of confocal and gSTED images of dendrite expressing Homer1c-mCherry and GFP-CLCa. Scale bars dendrite: 2 µm, zooms: 500 nm. (C) Number of clathrin-coated structures per PSD per neuron, represented as mean ± SEM (N = 12 neurons). (D) Scatterplot of the length (µm) and width (µm) of clathrin-coated structures in the dendritic shaft and associated with Homer1c based on ferret dimensions (spine: n = 248, shaft: n = 301). (E) Circularity ratio plotted against area (µm2) (spine: n = 248, shaft: n = 301). (F) Violin plot of clathrin area (µm2) in dendritic shaft and spines (spine: n = 248, shaft: n = 301).

Next, we analyzed the morphology of dendritic clathrin structures and found a large range of sizes from as small as 0.01 µm^2^ up to 0.43 µm^2^ (Figure 1E, F). On average, the area of PSD-associated clathrin structures was lower, albeit not statistically different from the average area of clathrin structures found in the shaft (area clathrin structures in shaft: 0.045 ± 0.003 µm^2^, spine: 0.038 µm^2^ ± 0.002, P > 0.1) (Figure 1F). However, the variability in sizes of clathrin structures in the shaft, was much larger than in spines (CV shaft: 1.3; spines: 0.84), with larger structures exclusively found in the shaft and not in spines (Figure 1D, E; range in width/length shaft: 0.15 µm to 1.7 µm, spine: 0.084 µm to 0.78 µm). Indeed, large clathrin structures were regularly observed in the shaft, approximately ∼3 per 20 µm of dendrite (data not shown). Thus, dendrites contain a large variation of clathrin-marked structures, with PSD-associated EZs being a distinct, homogenous sub-population of clathrin structures in dendritic spines.

Studies in non-neuronal cells often classify clathrin structures as flat lattices based on size and shape. Compared to small, circular clathrin structures, presumably representing endocytic pits or intracellular vesicles, lattices are defined as large and irregularly shaped clathrin structures (Grove et al., 2014; Leyton-Puig et al., 2017; Saffarian et al., 2009). We generated scatterplots of the circularity and area of individual dendritic clathrin structures to test if we could find a similar classification (Figure 1E). However, there was no correlation between size and circularity in dendritic clathrin structures (R2 = 0.006). Also, we did not observe a clear differentiation in clathrin structures using this approach, suggesting that clathrin-coated structures in dendrites form a highly heterogeneous population that cannot be classified based on these morphological parameters. Altogether, these data highlight the morphological heterogeneity of clathrin-coated structures in neuronal dendrites

### Clathrin dynamics at the EZ are distinct from clathrin-coated structures in the dendritic shaft

To study the dynamic properties of clathrin-coated structures in both spines and dendritic shaft we next performed live-cell imaging of GFP-CLCa in dendrites. We first investigated the dynamics of clathrin-coated structures on short time intervals by imaging at 0.2 Hz for 5 minutes. To differentiate stationary from moving particles we used a Fourier analysis-based filtering on kymographs (Mangeol et al., 2016). The dendritic shaft predominantly contained stationary clathrin-coated structures, however smaller anterograde and retrograde moving puncta were also observed (Figure 2A). Interestingly, these fast-moving particles were small (IQR: 0.013 – 0.065 µm^2^) most likely reflecting intracellular vesicle transport. Within the stationary pool in the shaft, we observed a few distinct clathrin-coated structures. The two most frequently observed structures were larger, high-intensity structures that either remained fluorescently stable over the entire course of imaging (CV fluorescence intensity = 0.09, Figure 2B: upper panel), or showed large fluctuations in fluorescence intensity (CV = 0.16, Figure 2B: lower panel), perhaps indicating a more dynamic structure. In rare cases transient budding of clathrin from these structures was observed, reminiscent of endocytic pit formation.

**Figure 2.**
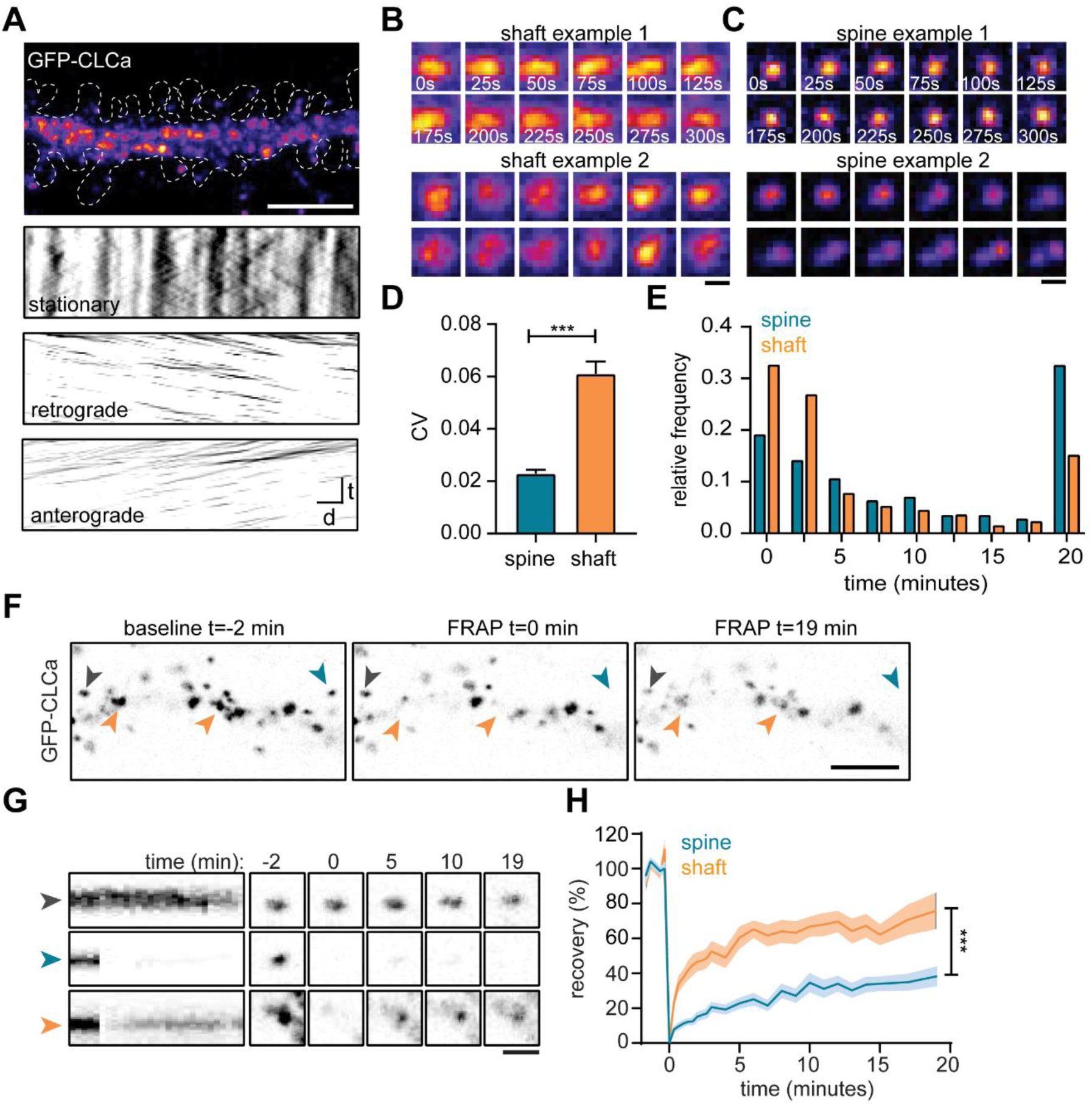
The EZ is dynamically distinct from shaft clathrin-coated structures. (A) Representative dendrite expressing GFP-CLCa, scale bar: 5 µm, and kymographs of clathrin-coated structures in the dendritic shaft only, separated in stationary (upper panel), retrograde (middle panel) and anterograde (lower panel) particles. Scale: time t, on the y-axis is 5 minutes, and distance d on x-axis is 20 µm. (B) Two examples of intensity fluctuations in stationary dendritic shaft structures. Scale bar: 1 µm. (C) Two examples of intensity fluctuations in spine structures. Scale bar: 1 µm. (D) Fluctuations in intensity plotted as the coefficient of variance (CV) between shaft and spine (spine: n = 48, shaft: n = 49, p < 0.001). Data represented at mean ± SEM. (E) Histogram of the lifetime of clathrin-coated structures in shaft and spine (spine: n = 171, shaft n = 769), data represented as fraction. (F) Example images of GFP-CLCa before (left panel), directly after FRAP (middle panel) and recovery (right panel), scale bar: 5 µm. Grey arrow indicates control, unbleached region, blue indicated bleached EZ, orange indicates bleached stationary dendritic shaft structures. (G) Kymograph and example images of the structures indicated in F. Kymograph shows 22-minute acquisition, scale bar: 1 µm. (H) Percentage of recovery in shaft (orange, n = 14) and spine (blue, n= 30).

In spines, the EZ appeared much more stable than dendritic clathrin structures, with little fluctuations in GFP-CLCa intensity (CV: 0.02, Figure 2C: upper panel). Strikingly, we were able to pick up, what seemed to be the budding of individual vesicles from the EZ (Figure 2C, lower panel). On average, the fluctuations in intensity of clathrin structures in shaft and spines were significantly different, with much lower fluctuations found in spines (shaft CV: 0.06 ± 0.004, spine CV: 0.02 ± 0.001, p < 0.001) (Figure 2D). Longer acquisitions of 20 minutes at 30-second intervals showed that the clathrin-coated structures in spines had a considerably higher average lifetime compared to clathrin-coated structures in the shaft (average lifetime spines: 10 ± 0.7 minutes, shaft: 6.0 ± 0.3 minutes, p < 0.001) (Figure 2E). Indeed, 68.1 ± 6.0% of PSDs remained associated with at least one clathrin structure that was present for the entire 20 minutes, confirming that the EZ is stably coupled to the PSD (Blanpied et al., 2002; Lu et al., 2007; Scheefhals et al., 2019). In contrast, in the dendritic shaft, only a small fraction (∼15%) of clathrin-coated structures was long-lived (>17.5 minutes) and the median lifetime of all events was ∼2.5 minutes, indicating that the majority of clathrin structures in the dendritic shaft are transient structures (Figure 2E).

The relatively long lifetime and small fluctuations in intensity of clathrin at the EZ might suggest a considerably lower turnover of clathrin at the EZ compared to shaft structures. To determine the turnover of clathrin at stable dendritic structures we used fluorescence recovery after photobleaching (FRAP) of GFP-CLCa (Figure 2F). A relatively long baseline of two minutes was acquired to make sure that only stationary structures would be included in the analysis. We determined that in stable shaft structures GFP-CLCa recovered relatively fast (tau: 13.0 min) to 75.8 ± 10% in 20 minutes (Figure 2G, H), indicating a high level of clathrin exchange at these structures. In contrast to the high turnover of stationary structures in the shaft, the EZ showed relatively low levels of turnover (tau: 36.2 min) and total recovery (38.2 ± 5.2% after 20 minutes) (Figure 2G, H), suggesting little exchange of clathrin at the EZ. Taken together, these live-cell imaging experiments show that clathrin-coated structures in the dendritic shaft are morphologically and dynamically highly diverse, and that the EZ in dendritic spines contains a stable accumulation of clathrin that is very similar from spine to spine, thereby differentiating itself from all other clathrin-coated structures.

### Nanoscale organization of the endocytic zone in dendritic spines

To further resolve the spatial organization of the EZ, we next used single-molecule localization microscopy (SMLM). Homer1c-mCherry and GFP-CLCa were co-transfected as before and labelled with primary and secondary antibodies to perform two-color dSTORM imaging and reconstruct high-density localization maps of the distribution of clathrin molecules within the EZ and relative to the PSD (Figure 3A). We used DBScan to define clusters of Homer1c molecules, outlining the PSD and the associated clathrin clusters, marking the EZ (Figure 3B). We found that the centroid of the EZ was generally located within 100 nm from the border of the PSD (Figure 3C) with an average border-to-centroid distance from PSD to EZ of 10.6 ± 7.8 nm, confirming that the EZ is closely linked to the PSD and well within a distance that can be linked by scaffold proteins (Lu et al., 2007).

**Figure 3.**
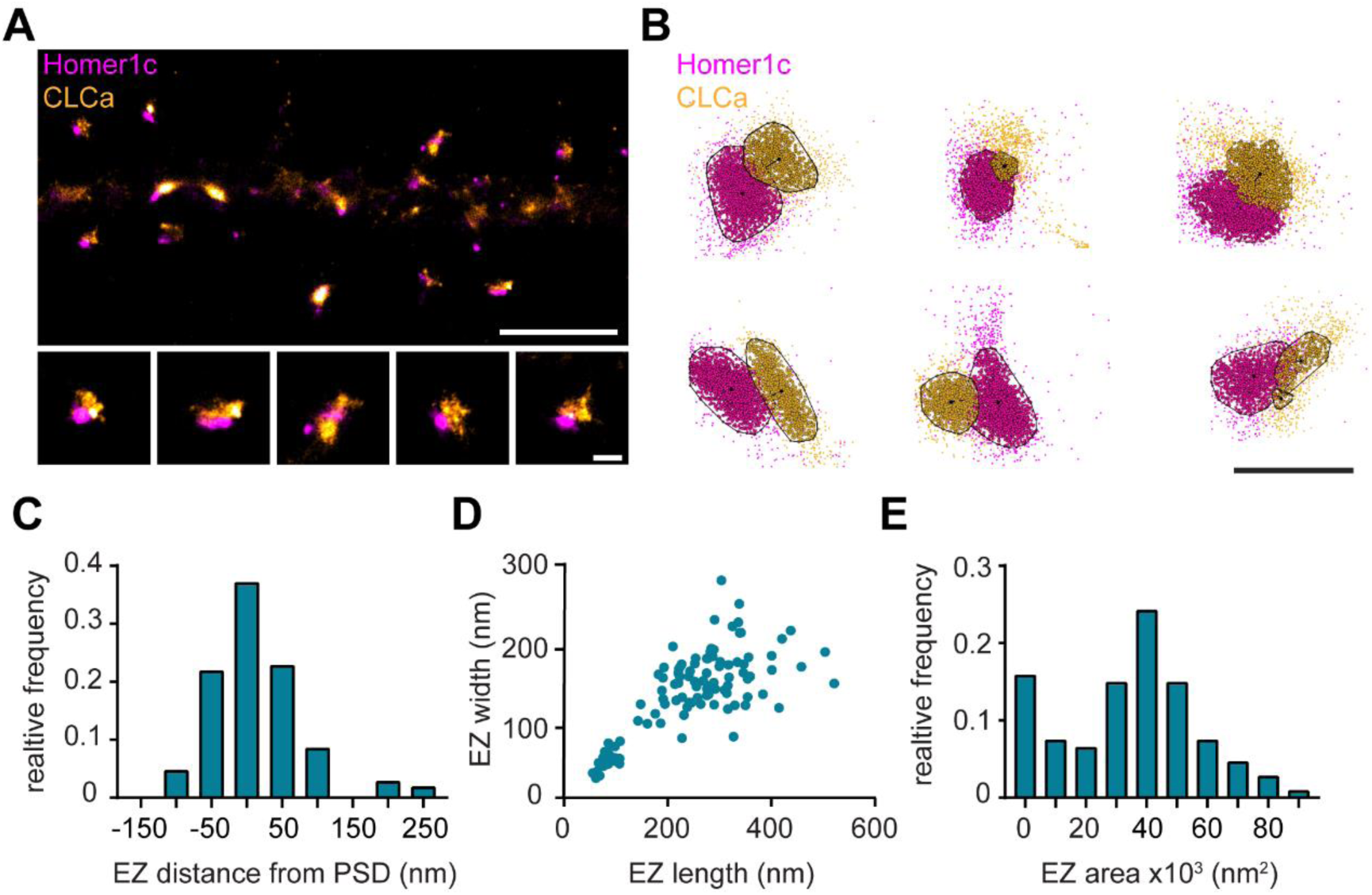
Nanoscale organization of the endocytic zone. (A) SMLM image of dendrite expressing Homer1c-mCherry and GFP-CLCa labelled with CF568 and A647 respectively and zooms of individual EZs. Scale bar upper panel: 2 µm, zooms: 250 nm. (B) Individual molecules of Homer1c (magenta) and CLCa (orange) are outlined using DBScan. Black dot and line indicate center of the EZ (dot) and distance to the border of the PSD (line). Scale bar: 500 nm. (C) Histogram of the border (Homer1c) to center (CLCa) distance in nm. (D) Scatterplot of the FWTM length (nm) and FWTM width (nm) of the EZ. (E) Histogram of the area of the EZ plotted as x10^3^ nm^2^. (C-E) n = 107.

On average the area of the EZ was 35.0 ± 2.0 x10^3^ nm^2^ (Figure 3E), and 224.6 ± 10.4 nm in length and 146.5 ± 5.2 nm in width (Figure 3D). Moreover, the dimensions (length and width) of individual structures were positively correlated (R2 = 0.58). Interestingly, we often found that PSDs were associated with multiple clathrin structures, similar as to what we observed with gSTED imaging (Figure 1B, D). We noted that two distinct populations could be observed based on morphological characteristics and distinguished between the primary and secondary clathrin structure based on size. The largest was classified as the primary structure and we found that this structure most likely corresponds to the EZ, as these were also the most closely linked to the PSD (Supplement Figure 2A, B). The secondary, smaller structures were between 50 and 100 nm in diameter (Supplement Figure 2C), similar to the reported size of endocytic vesicles (Kirchhausen and Harrison, 1981; Pearse and Crowther, 1987). These smaller structures also appeared more circular and further away from the PSD (Supplement figure 2B, D), further suggesting that these smaller secondary structures are endocytic vesicles that perhaps budded off from the edge of the EZ.

### Endocytic proteins are differentially retained at perisynaptic sites

Apart from clathrin, only a few other endocytic components have been suggested to be part of the EZ. Among these proteins are synaptotagmin-3 (Awasthi et al., 2018), PICK1 (Maria Fiuza et al., 2017) and CPG2 (Cottrell et al., 2004). Moreover, the presence of dynamin2 and AP2 at the site of clathrin-coated pits in spines suggest that these proteins could also be part of the EZ (Rácz et al., 2004). However, it remains unknown whether these and other endocytic proteins are stably accumulated at the EZ, or whether these are perhaps transiently recruited only during endocytic events. To begin to address this, we first determined the localization of 12 well-known endocytic proteins using confocal microscopy and live-cell imaging. Among these proteins are the well-known F-BAR and N-BAR proteins like FCHO1, syndapin-1 (Sdp1), syndapin-2 (Sdp2) and amphiphysin (Amph), β2-adaptin, a subunit of the membrane proteins AP2, scission protein dynamin2 (Dyn2) and other adaptor proteins like Eps15, PICALM, intersectin-1 long (Itsn1L) and epsin-2 (Epsn2). In addition, we included HIP1R and CPG2 that can couple the endocytic machinery to the actin cytoskeleton (Chen and Brodsky, 2005; Engqvist-Goldstein et al., 2001; Loebrich et al., 2016; Wilbur et al., 2008). We found that most of these proteins localized at perisynaptic sites in a punctate manner, similar to clathrin (Figure 4B). Sdp1, Sdp2 and Amph showed a more diffuse signal within the spine and dendritic shaft. For Amph, clear puncta associated with the PSD could be detected occasionally, however Sdp1 and Sdp2 did not seem to be enriched in distinct puncta and were not further analyzed. In this experiment we found that 66.8 ± 6.9% of PSDs was associated with GFP-CLCa (Figure 4A, C). We found that the fraction of PSDs associated with HIP1R, β2-adaptin, Dyn2, CPG2, Eps15, and Itsn1L was similar to the percentage of clathrin-associated PSDs. In contrast, PICALM, Epsn2, Amph and FCHO1 were less frequently found in association with the PSD (Figure 4B, C). Thus, HIP1R, β2-adaptin, Dyn2, CPG2, Eps15 and Itsn1L appear associated with the PSD and could be intrinsic components of the EZ.

**Figure 4.**
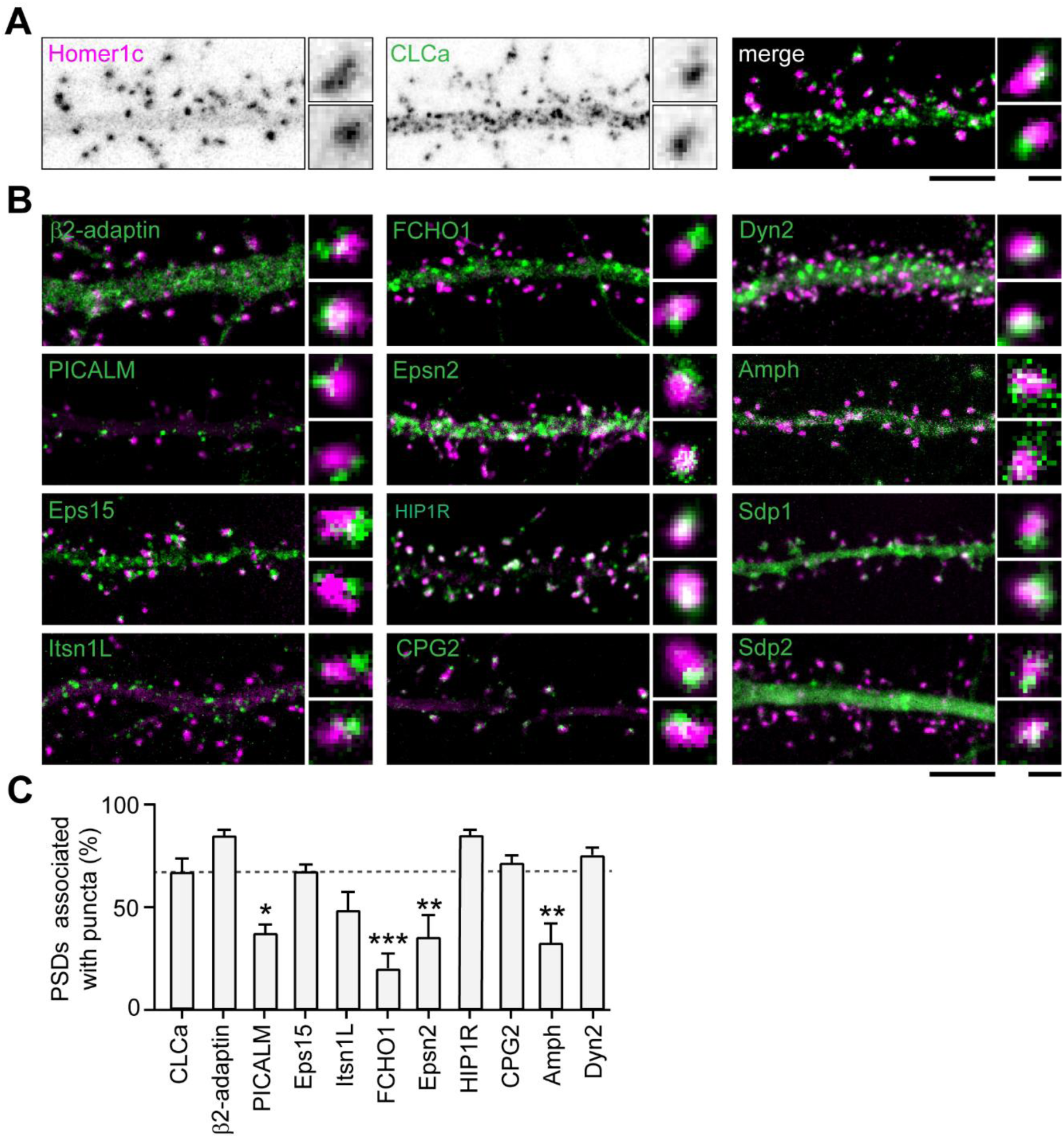
Endocytic accessory proteins localize to the perisynapse. (A) Example images of dendrites expressing Homer1c-mCherry and GFP-CLCa visualized as black and white images (left, middle panel), and merge (right panel). (B) Example images of neurons co-expressing tagged endocytic proteins relative to Homer1c. (A-B) Scale bars: 5 µm, zoom dimensions: 1 µm. (C) Percentage of synapses associated with endocytic proteins, represented as mean ± SEM. Relative to Homer1c-CLCa association (n = 8), PICALM-mCherry (n = 6, p < 0.05), FCHO1-mCherry (n = 6, p < 0.001), Epsn2-mCherry (n = 9, p < 0.01) and Amph-mCherry (n = 6, p < 0.01) were significantly less often associated with the PSD, while β2-adaptin-GFP (n = 6), GFP-Eps15 (n = 5), GFP-Itsn1L (n = 5), HIP1R-GFP (n = 9), GFP-CPG2 (n = 5), Dyn2-GFP (n = 5) were not different from GFP-CLCa. Data represented as mean ± SEM.

Next, to test whether these endocytic proteins were stably associated with the PSD we performed time-lapse experiments on neurons co-expressing Homer1c and a fluorophore-tagged endocytic protein. Neurons were imaged for 10 minutes at 20-second time intervals. We found very distinct behaviors in the dynamics of endocytic proteins. While some proteins only transiently occurred at perisynaptic sites (e.g., FCHO1; Figure 5A), other proteins appeared stable over the entire duration of the acquisition (e.g., CPG2; Figure 5B). Consistent with our previous observations in fixed neurons (Figure 4C), the percentage of PSDs associated with a clear endocytic protein structure for the entire duration of the acquisition was high for HIP1R, β2-adaptin, Dyn2, CPG2, Eps15 and Itsn1L, and much lower for PICALM, FCHO1, Epsn2 and Amph (Figure 5C). From these live-cell acquisitions we determined the lifetime of events where these proteins were enriched at perisynaptic sites. Interestingly, when plotted as histograms, a clear bimodal distribution of lifetimes was observed (Figure 5D). Endocytic proteins accumulated either briefly (<3 min) or appeared persistent (>9 minutes) at perisynaptic sites (Figure 5D). These data also indicated that HIP1R, β2-adaptin, Dyn2, CPG2, Eps15 and Itsn1L are stable components that are generally long-lived (Figure 5D). In contrast, the average lifetimes of FCHO1 (2.74 ± 1.7 min), PICALM (1.73 ± 0.2 min), Epsn2 (3.00. ± 0.2 min), and Amph (2.73 ± 1.7 min) at perisynaptic sites were significantly lower compared to the average lifetime of CLCa. Notably, the lifetime of these short-lived events is comparable to the duration of endocytic events (∼2 minutes), suggesting that these proteins are transiently recruited upon the induction of endocytosis. To determine the turnover of the long-lived endocytic proteins at perisynaptic sites we performed FRAP experiments. Except for CPG2, all endocytic proteins showed considerably higher turnover than clathrin (percentage of recovery after 10 minutes CPG2: 29.4 ± 3.7%, HIP1R: 59.2 ± 5.2%, β2-adaptin: 56.7 ± 3.8%, Eps15: 88.1 ± 3.8, Itsn1L: 93.9 ± 8.7) (Figure 5E, Supplement Figure 3A-F). Taken together, these experiments reveal that apart from clathrin, HIP1R, β2-adaptin, Dyn2, CPG2, Eps15, and Itsn1L are also integral components of the perisynaptic EZ, while PICALM, FCHO1, Epsn2 and Amph only appear transiently at perisynaptic sites, perhaps to initiate or facilitate endocytosis.

**Figure 5.**
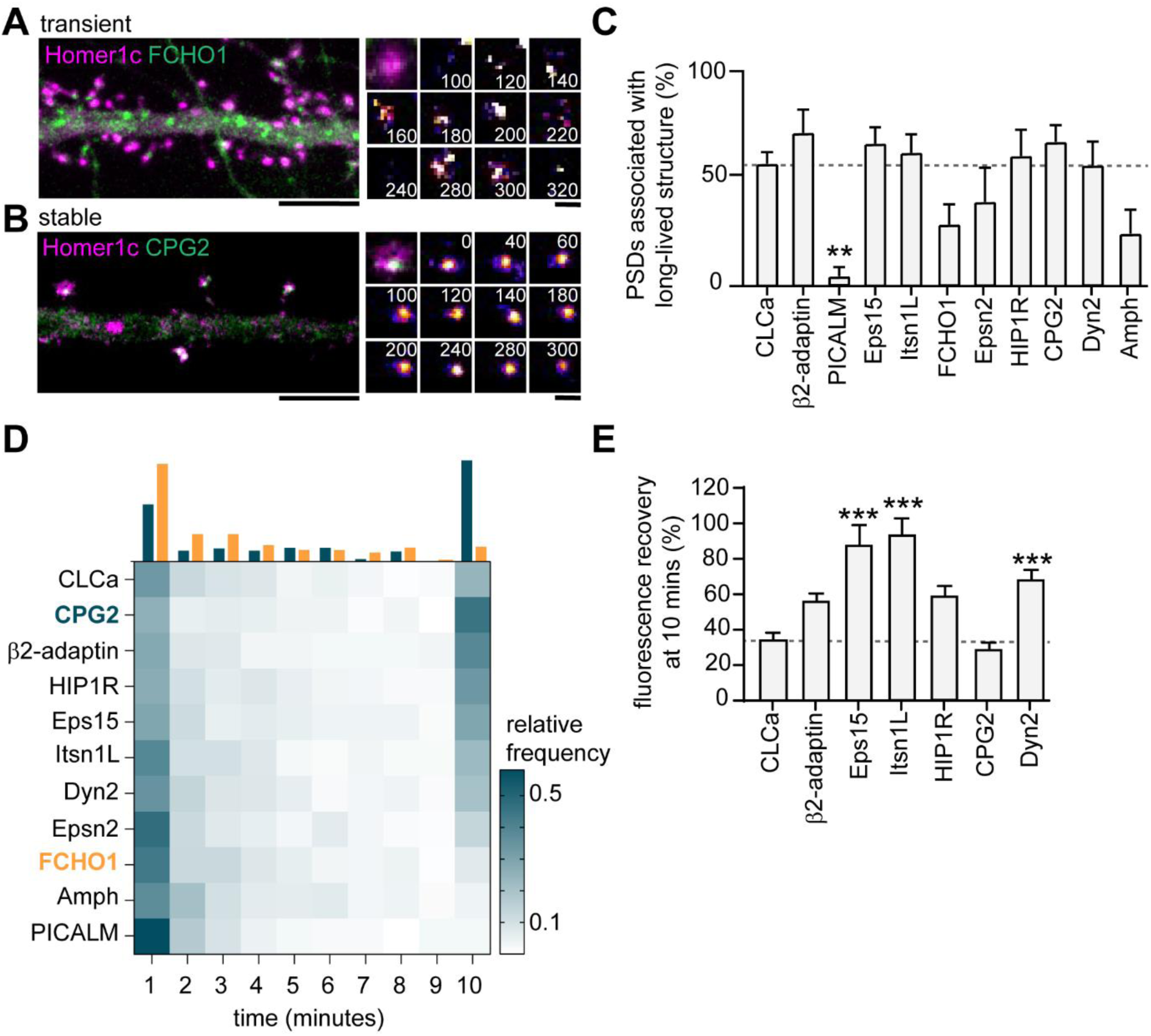
Endocytic proteins are differentially associated with the PSD. (A) Example image of FCHO1 (green) that is transiently associated with Homer1c (magenta). Zooms show temporal recruitment of FCHO1. Scale bar: 5 µm, zoom: 500 nm. (B) Example image of CPG2 that is stably associated with Homer1c. Zooms show temporal dynamics of CPG2. Scale bar: 5 µm, zoom: 500 nm. (C) Percentage of synapses that contain at least one stable structure (persisting for >9 minutes). Only PICALM-mCherry (n = 5, p < 0.01) was significantly less often stably associated with the PSD compared to GFP-CLCa (N = 6). β2-adaptin-GFP (N = 6), GFP-Eps15 (N = 6), GFP-Itsn1L (N = 6), FCHO1-mCherry (N = 5), Epsn2-mCherry (N = 5), HIP1R-GFP (N = 6), GFP-CPG2 (N = 8), Amph-mCherry (N = 5), Dyn2-GFP (N = 7), were not different from GFP-CLCa. (D) Heatmap visualizing the frequency distribution of the lifetime of endocytic proteins associated with the PSD. The histogram on top is an example of FCHO1 (orange) that is mostly short-lived, and CPG2 (blue) that is mostly stable, plotted as relative frequency. (E) Summary graph of the recovery 10 minutes after FRAP for GFP-Eps15 (n = 23 , p < 0.001), GFP-Itsn1L (n = 20, p < 0.001), HIP1R-GFP (n = 44, p < 0.01), Dyn2-GFP (n = 51, p < 0.001) had significantly higher turnover compared to GFP-CLCa (n = 32). GFP-CPG2 (n = 22) and β2-adaptin GFP (n = 13) were not different compared to GFP-CLCa. Data plotted as mean ± SEM.

### Endocytic proteins have distinct spatial organization relative to the clathrin coat at the EZ

The presence of multiple endocytic adaptor proteins and their stable retention at perisynaptic sites suggests that these proteins might be an integral part of the EZ. First, we applied two-color gSTED on Halo-CLCa co-transfected with the stably retained endocytic proteins fused to GFP and stained for endogenous Homer1b/c to localize the PSD. We indeed found that Eps15, Itsn1L, Dyn2, β2-adaptin and HIP1R all colocalize with clathrin next to the PSD (Supplement Figure 4A-C).

Next, to dissect the spatial organization of endocytic proteins relative to the clathrin structure at the EZ we used two-color SMLM. Halo-CLCa was co-transfected with a GFP-tagged endocytic protein to efficiently label and acquire high-density localization maps in two channels. Strikingly, we found that β2-adaptin, Eps15, and Itsn1L were often distributed in smaller patches around and sometimes within the EZ marked by CLCa (Figure 6A-C). HIP1R showed a more homogenous distribution and often colocalized with the EZ entirely and even surrounding the EZ (Figure 6D). Dyn2 showed an overall more homogenous distribution, similar to HIP1R (Figure 6E). However, we also found examples where Dyn2 localized in small clusters at the edge of the EZ. To analyze these distributions quantitatively, we manually selected regions around clathrin structures in dendritic spines for further analysis. We then used DBScan to determine the outline of the EZ marked by CLCa. We first mapped the absolute distance of localizations to the border of the EZ averaged over a population of EZs and found that all the endocytic proteins analyzed here were preferentially localized at the edge of the clathrin structure (Figure 6F). However, since individual EZs can vary in size, we next mapped the density of endocytic proteins in rings that were set in size relative to the clathrin structure. For each EZ, we defined 8 incremental rings that were scaled with proportion to the outline of the EZ and binned the density of localizations within each of these rings (Figure 6G). As expected, when plotting the relative fraction of Halo-CLCa localization within the rings, we found that the density of clathrin molecules was highest in the center (0-20% ring), gradually decreased towards the outer ring (80-100%) and dropped to close to zero in the rings surrounding the EZ (100-160% rings). In contrast, when we plotted the relative density of Homer1c localizations relative to CLCa, we found a clear separation of these distributions (Supplement Figure 2E, F), further validating the analysis. When analyzing the endocytic adaptor proteins relative to the EZ, we again found that the relative density of Eps15, β2-adaptin and Itsn1L peaked within the EZ, but close to the edge of the EZ (Figure 6G). The profiles of HIP1R and Dyn2 localizations (Figure 6G) showed less clear peaks, indicating a more homogenous distribution of these proteins within the EZ.

**Figure 6:**
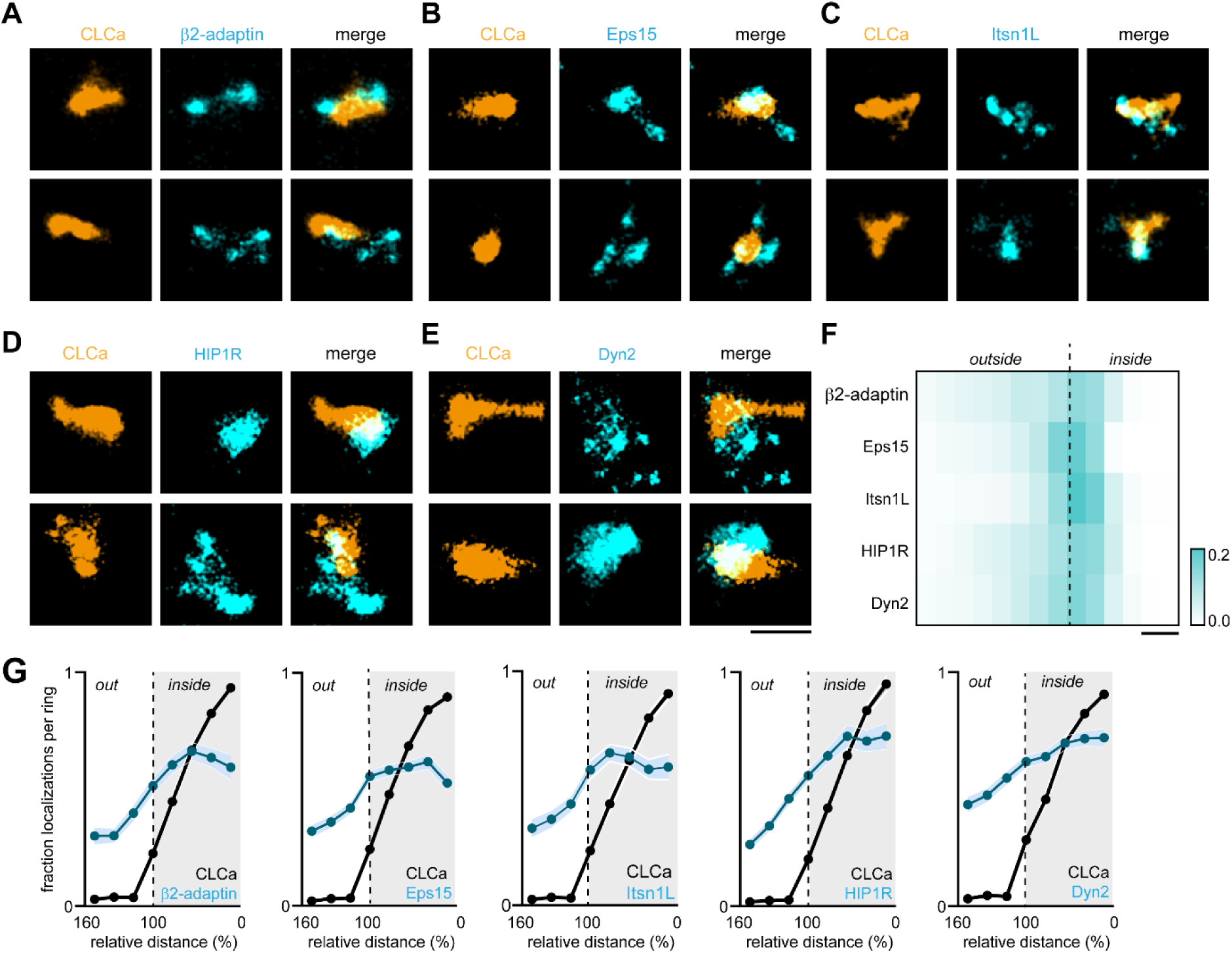
Endocytic proteins have distinct spatial organization relative to the clathrin structure marking the EZ. (A-E) High resolution example images of Halo-CLCa labelled with JF647 (orange) co-expressed with endocytic proteins fused to GFP labelled with CF568 (cyan). Scale bar: 500 nm. (F) Heatmap visualizing the relative frequency of the distance of individual localizations relative to the border of CLCa. (G) Fraction of localizations per ring. Dotted line depicts the border of CLCa. β2-adaptin (n = 87), GFP-Eps15 (n = 126), GFP-Itsn1L (n = 58), GFP-HIP1R (n = 72), GFP-Dyn2 (n = 82).

### Interactions with the PSD, but not with the membrane or actin cytoskeleton are required for the perisynaptic localization of the EZ

The differential dynamics and nanoscale organization of endocytic proteins at perisynaptic clathrin structures suggests that the EZ is a highly organized structure where several endocytic proteins are assembled. The mechanisms that retain the EZ at this particular position, however, are not fully understood. The EZ is coupled to the PSD via Shank-Homer-Dyn3 interactions (Lu et al., 2007; Petrini et al., 2009; Scheefhals et al., 2019). In addition, we now identified several new EZ components that can couple to the plasma membrane, e.g., via AP2, or the actin cytoskeleton, e.g., via CPG2 and HIP1R (Loebrich et al., 2016; Saffarian et al., 2009), suggesting that these modes of interaction could also contribute to the positioning of the EZ. To test this, we interfered with several of these connections. First, we tested whether Shank knockdown (KD), which we showed previously uncouples the EZ from the PSD (Scheefhals et al., 2019), also leads to the loss of these newly identified EZ components (Figure 7A, B). Indeed, we found that Shank-KD did not only reduce the number of clathrin-positive PSDs as found before (0.5 ± 0.1 relative to control), but also reduced the association of the PSD with other endocytic proteins (Figure 7B). We found that Shank-KD significantly reduced Homer1c-Eps15 (0.63 ± 0.1), Homer1c-Itsn1L (0.6 ± 0.1) and Homer1c-β2-adaptin (0.55 ± 0.1) coupling compared to control (Figure 7B). Interestingly, HIP1R (0.89 ± 0.04) and Dyn2 (0.91 ± 0.1) were not uncoupled from the PSD (Figure 7B). Together, these findings show that PSD-EZ coupling via Shank proteins is necessary for the integrity of the EZ. Next, to test if other mechanisms contribute to EZ maintenance we tested whether alterations in actin dynamics, membrane binding capacity or depleting specific EZ components would lead to a reduction of EZs (Figure 7C). Interestingly, we found that disrupting the integrity of the actin cytoskeleton did not result in a clear reduction of the PSD-EZ association. The actin depolymerization drug Latrunculin B slightly decreased the PSD-EZ association (0.81 ± 0.1), but this was not statistically significant. The Arp2/3 inhibitor CK-666 (0.98 ± 0.1) also did not significantly alter PSD-EZ association (Figure 7D, F), suggesting that disrupting actin dynamics does not disrupt positioning of the EZ. Jasplakinolide, an F-actin stabilizing drug, resulted in significantly more PSD-EZ association compared to control (1.27 ± 0.04, p<0.05). To test specific EZ-actin interactions, we assessed whether the binding of clathrin to HIP1R is necessary for EZ maintenance. Overexpression of a clathrin-light chain mutant that is unable to bind HIP1R (GFP-CLCb-EED/QQN) (Chen and Brodsky, 2005; Poupon et al., 2008), did not affect the localization of the EZ (Figure 7E, F), further indicating that coupling to the actin cytoskeleton is not a primary mechanism for maintaining the EZ. The third mechanism that could allow the perisynaptic localization of the EZ involves interactions with the plasma membrane. To address this, we overexpressed an AP2m1 mutant that is unable to interact with PIP2 and was shown to hamper receptor internalization (Raman et al., 2014), but we found no change in the fraction of EZ-positive PSDs (1.1 ± 0.1), suggesting that coupling to the membrane via AP2 does not affect EZ maintenance (Figure 7E, F). Lastly, we checked if removing a specific, stable endocytic component, Itsn1L would affect the EZ. Itsn1L is multi-domain scaffold protein that interacts with several endocytic proteins to orchestrate endocytosis and could thus have a central role as scaffold in the EZ. However, Itsn1L knockdown did not alter the fraction of EZ-positive PSDs (Figure 7E, F). Altogether, based on these mechanistic experiments, we conclude that the EZ is assembled from a distinct set of endocytic proteins and is maintained and positioned primarily by interactions with the PSD.

**Figure 7.**
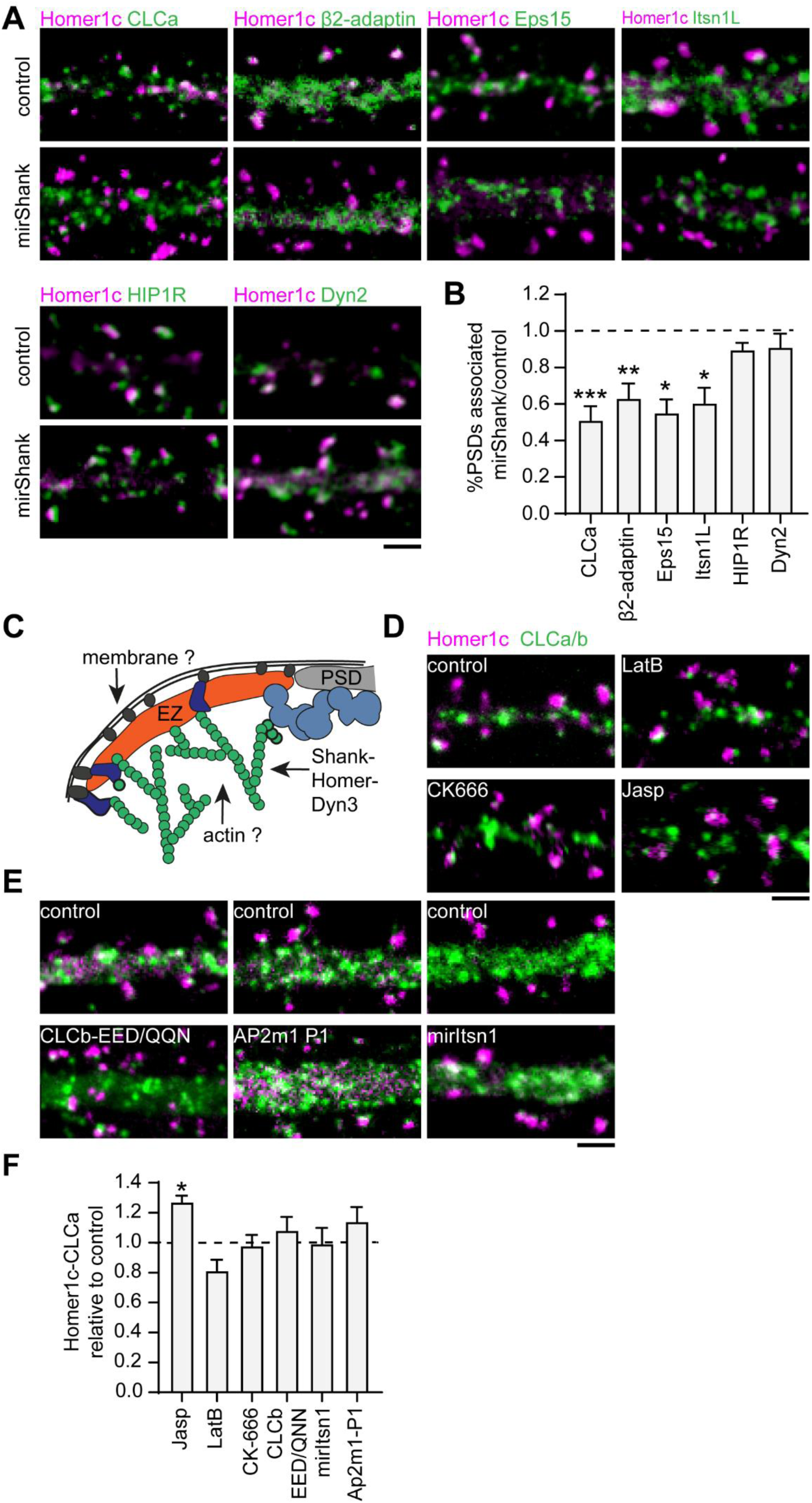
Interactions with the PSD, but not with the membrane or actin cytoskeleton are required for positioning of the EZ. (A) Example images of dendrites expressing Homer1c-ALFA and endocytic proteins fused to GFP co-expressed with control or mirShank-mCherry construct. Scale bar: 2 µm. (B) Fraction of PSDs associated with an EZ after Shank-KD relative to control plotted as mean ± SEM. GFP-CLCa (N = 8, p < 0.001), β2-adaptin (N = 10, p < 0.01), Eps15 (N = 12, p < 0.05), Itsn1L (N = 10, p < 0.05), HIP1R (N = 11 p >0.05), Dyn2 (n = 12, p > 0.05). (C) Illustration of possible mechanisms that could maintain the EZ adjacent to the PSD. (D) Example images of dendrites co-expressing Homer1c-ALFA and GFP-CLCa in dendrites treated with LatB, CK666 or Jasp. Scale bar: 2 µm. (E) Example images of dendrites expressing control constructs, CLCb-EED/QQN (left panel) and AP2m1-P1 (middle panel), or Itsn1 KD construct (mirItsn1; right panel). Scale bar: 2 µm. (F) Fraction of EZ-associated PSDs relative to control, plotted as mean ± SEM. Jasp (N = 4), LatB (N = 8), CK666 (N = 12), CLCb-EED/QQN (N = 7), mirItsn1 (N = 11), AP2m1-P1 (N = 5).

## DISCUSSION

Neurons contain a large variety of clathrin structures. A particularly important clathrin structure in neurons, the postsynaptic EZ, is characterized by the stable accumulation of clathrin associated with the PSD. Localized endocytosis of synaptic receptors at the EZ is essential for the maintenance and activity-directed changes in the composition of the synaptic membrane. However, despite vigorous investigation of clathrin-coated structures in various cell types, the molecular composition and organization of the EZ has remained largely elusive. Here, we present evidence that the EZ is a highly unique clathrin structure. We found that a defined arsenal of endocytic proteins is differentially retained at the EZ and highly organized at the nanoscale level with respect to the clathrin assembly.

Our data show that the postsynaptic EZ can be clearly distinguished from other dendritic clathrin assemblies. Clathrin-coated structures in the dendritic shaft form a highly heterogenous population, including small, fast-moving particles, as well as larger stationary structures. This heterogeneity in clathrin-coated structures resembles those found in other cell types, where a large variety of clathrin assemblies have been identified. Often, small clathrin structures represent transient endocytic pits or intracellular vesicles while the large patches are stable, membrane-attached structures (Grove et al., 2014; Leyton-Puig et al., 2017; Saffarian et al., 2009). Indeed, we found that the small structures in the dendritic shaft are often transient or moving, most likely representing endocytic pits or cargo vesicles, while the larger assemblies are stationary but still undergo dynamic exchange of clathrin. The function of these larger dendritic patches remains unknown. In dendritic spines, we found that the morphological characteristics of the EZ are highly similar from spine to spine, with much less variation in size and dynamics than observed for clathrin structures in the dendritic shaft. Moreover, the EZ appeared as a long-lived and fluorescently stable clathrin structure with relatively low exchange of clathrin, in line with previous studies (Blanpied et al., 2002; Petrini et al., 2009; Rosendale et al., 2017; Scheefhals et al., 2019). Together, these results indicate that based on the morphological and dynamic behavior of clathrin, the EZ can be distinguished from other clathrin assemblies found in the dendritic shaft.

The EZ has been postulated as a primary site for endocytosis of synaptic membrane proteins to sort these components in the local recycling machinery and effectively retain these at the synaptic membrane (Blanpied et al., 2002). Indeed, several studies have unequivocally demonstrated that in the absence of the EZ endocytic trafficking of glutamate receptors is severely affected, leading to the net loss of membrane-expressed receptors and consequential deregulated glutamatergic signaling (Cortese et al., 2016; Lu et al., 2007; Nakano-Kobayashi et al., 2014; Petrini et al., 2009; Scheefhals et al., 2019). Formation of endocytic vesicles containing synaptic receptors has also been directly visualized in close proximity to the PSD (Rosendale et al., 2017), supporting the endocytic capacity of the EZ. In our live-cell experiments, we occasionally observed the formation of a secondary clathrin structure from the larger, primary clathrin structure. Further, super-resolution imaging allowed us to resolve the EZ at higher resolution and we often found that PSDs were associated with more than one clathrin structure, with the secondary, smaller structure often having a size similar to those described for clathrin-coated vesicles (Kirchhausen et al., 2014). These secondary structures were often 200 - 250 nm away from the border of the PSD, i.e., more distant from the PSD than the EZ that was closely (within ∼30 nm) associated. Similarly, EM studies showed that clathrin coated vesicles bud off from the membrane preferentially at 100 - 600 nm from the PSD (Rácz et al., 2004). Thus, although we did not reach sufficient resolution to unequivocally resolve clathrin-coated pits associated with the EZ, our data is consistent with the idea that endocytic pits bud off at the edge of the EZ.

Importantly, our findings significantly expand on the notion that the EZ is a perisynaptic site of endocytosis by identifying several key endocytic proteins that reside at the EZ. Based on our live-cell imaging, quantitative super-resolution imaging and mechanistic studies we conclude that β2-adaptin, Eps15, and Itsn1L are stable EZ residents that localize preferentially at the edge of the EZ. HIP1R and Dyn2 were also stably associated with the EZ, but this association seemed independent of EZ-PSD coupling. PICALM, FCHO1, Epsn2 and Amph were associated with the EZ to a much lesser extent and appeared only transiently at perisynaptic sites. Thus, different classes of endocytic proteins, both early- and late-phase proteins are associated with the EZ.

The early-phase endocytic proteins β2-adaptin, Epsn2, Eps15, Itsn1L and FCHO1 are all vital for the initial stages of endocytosis (Cocucci et al., 2012; Henne et al., 2010; Kirchhausen et al., 2014; Reider et al., 2009; Saffarian et al., 2009). We found that the localization patterns of these proteins in neurons were overall very similar, with clear puncta throughout the dendritic shaft and spines. However, in contrast to β2-adaptin, Eps15 and Itsn1L that were stably associated with the PSD, Epsn2 and FCHO1 appeared solely as transient, short-lived puncta close to the PSD. Interestingly, the FCHO and Epsin proteins contain F-BAR domains and preferentially bind curved membranes. Thus, perhaps these proteins are only transiently recruited to the EZ upon induction of endocytosis and the associated increase in membrane curvature. β2-adaptin, Eps15 and Itsn1L were stably associated with the PSD, and located preferentially at the edge of the clathrin structure, surprisingly consistent with findings on flat clathrin lattices in non-neuronal cells (Sochacki et al., 2017). The accumulation of the AP-2 complex and its binding partner Eps15 at the periphery of the EZ likely contributes to the efficient capture of cargoes, i.e., synaptic membrane proteins, and their local uptake via endocytosis. Itsn1 is a multi-domain scaffold protein that coordinates different aspects of endocytosis. Itsn1 interacts with AP-2 (Pechstein et al., 2010), acts as a GEF for Cdc42 (Hussain et al., 2001) , and regulates dynamin recruitment (Evergren et al., 2007). Thus, Itsn1 could have a central organizing role at the EZ. Consistently, we recently reported that Itsn1 knockdown abrogates mGluR-mediated AMPAR trafficking (van Gelder et al., 2020). The results presented here show that Itsn1 knockdown does not alter the location or overall morphology of the EZ indicating that while Itsn1 has likely an important role in coordinating endocytosis at the EZ, Itsn1 does not seem to directly support the maintenance of this clathrin structure.

The late-phase endocytic proteins Dyn2 and Amph are both required for the efficient fission of endocytic vesicles, the final step of endocytosis. Amph, like the other BAR proteins Epsn2 and FCHO1, appeared mostly diffusely distributed throughout the dendrite and was only transiently associated with the PSD. Dyn2 however, seemed stably retained close to the PSD and localized at the EZ. In most cases, we found that the distribution of Dyn2 at the EZ was diffuse, but the clear differential localization patterns between different EZs may indicate that dynamin relocates to the EZ upon endocytosis as has been suggested before (Rosendale et al., 2017). We found that the expression of the two Syndapins (or PACSINs), Sdp1 and Sdp2 was diffuse throughout the neuron and we found no clear association with the PSD. Nevertheless, Sdp1 was found to be involved in activity-induced internalization of glutamate receptors and sorting of vesicles after budding of the membrane (Anggono et al., 2013; Widagdo et al., 2016). Thus, the Syndapins seem not stably associated with the EZ, but could be involved in directing the endocytic vesicles that emerge from the EZ through the endosomal system.

The EZ is most likely associated with the membrane, however if and how this association is established remains unknown. Several proteins that we identified here as EZ components could couple the EZ to the membrane. For example, the AP2 complex directly binds clathrin heavy chain and phosphatidylinositol 4,5-biphosphate (PIP2) (Beacham et al., 2019; Gaidarov and Keen, 1999; Kadlecova et al., 2017; Mettlen et al., 2018; Owen et al., 2000; Shih et al., 1995; Traub et al., 1999) and we found that β2-adaptin was stably associated with the EZ. However, we found that expression of a dominant-negative form of AP2mu2 that cannot bind PIP2 did not abrogate EZ positioning. Another candidate, PICALM can also simultaneously bind PIP2 and clathrin (Ford et al., 2001), but we found that PICALM was only transiently associated with the PSD and is thus unlikely to form a stable intermediate between the EZ and the membrane. Thus, either these components are not involved in coupling the EZ to the membrane or membrane anchoring is not a prerequisite for EZ positioning.

The actin cytoskeleton is prominent in dendritic spines and supports many aspects of synaptic transmission and plasticity. Several endocytic proteins that we found enriched at the EZ could link the EZ to the actin cytoskeleton. For instance, both HIP1R and CPG2 are actin-binding proteins, were enriched in dendritic spines and associated with the PSD. Notably, these results are consistent with previous studies that found that CPG2 localizes to the EZ (Cottrell et al., 2004; Nedivi, 1999) and is essential for glutamate receptor trafficking (Loebrich et al., 2016; Loebrich et al., 2013). In fact, of all the proteins that we analyzed in the current study, CPG2 was the most persistently associated with the PSD, and appeared to be even more stable than clathrin, suggesting that CPG2 could have a function in stabilizing and maintaining the EZ. HIP1R directly binds clathrin light-chain and physically links clathrin to F-actin (Chen and Brodsky, 2005; Engqvist-Goldstein et al., 2001; Wilbur et al., 2008), to promote actin polymerization at endocytic sites. Moreover, HIP1R has been shown before to be involved in maintaining stable clathrin structures (Grove et al., 2014; Saffarian et al., 2009). Surprisingly however, neither disruption of the actin cytoskeleton with pharmacological compounds, nor interfering with the HIP1R-CLC association significantly disrupted the localization or integrity of the EZ. Thus, the actin cytoskeleton is most likely involved in facilitating endocytosis at the EZ, but does not seem to have a prime structural role in maintaining or positioning the EZ.

We and others have consistently found that the EZ is stabilized and associated with the PSD by Shank-Homer-Dynamin3 interactions (Lu et al., 2007; Scheefhals et al., 2019). Interestingly, we found that interfering with this interaction by depletion of Shank proteins, specifically the stable, early endocytic proteins, β2-adaptin, Eps15 and Itsn1L uncoupled from the PSD. On the other hand, HIP1R and Dyn2 remained enriched in spines after Shank knockdown. Together with our findings that HIP1R and Dyn2 are more homogenously distributed relative to the EZ suggests that these proteins are not directly coupled to the EZ under basal conditions but rather preferentially reside in close proximity via other interactions.

Taken all together, we found that the EZ is a highly organized clathrin structure where endocytic proteins are differentially retained and stabilized. This distinct organization likely facilitates the efficient capture and endocytosis of synaptic membrane proteins close to the PSD. These findings motivate further investigation into the molecular composition, the mechanisms that control the recruitment and activation of individual EZ components and the coupling of the EZ to the intracellular endosomal system. Elucidating these aspects of the EZ will contribute to a better understanding of this subcellular structure in neurons that is so critical for the maintenance and activity-dependent modulation of neuronal synapses.

## MATERIALS AND METHODS

### Animals

All animal experiments were performed in compliance with the guidelines for the welfare of experimental animals issued by the Government of the Netherlands (Wet op de Dierproeven, 1996) and European regulations (Guideline 86/609/EEC). All animal experiments were approved by the Dutch Animal Experiments Review Committee (Dier Experimenten Commissie; DEC), performed in line with the institutional guidelines of Utrecht University.

### Primary hippocampal cultures and transfection

Hippocampal cultures were prepared from brain of embryonic day 18 (E18) Wistar rats (both genders) as described before (Scheefhals et al., 2019). Dissociated hippocampal neurons were plated on coverslips coated with poly-L-lysine (37.5 µg/ml, Sigma-Aldrich) and laminin (1.25 µg/ml, Roche Diagnostics) at a density of 100,000 neurons per well of a 12-well plate. Cultures were allowed to settle in Neurobasal medium (NB) supplemented with 2% B27 (GIBCO), 0.5 mM glutamine (GIBCO), 15.6 mM glutamate (Sigma-Aldrich), and 1% penicillin/streptomycin at 37°C in 5% CO2. After 24 hours halve of the NB medium was refreshed with BrainPhys medium (BP) supplemented with SM1 supplement (Stemcell Technologies) and 1% penicillin/streptomycin, and kept at 37°C in 5% CO2. Refreshment were done weekly replacing halve of the medium with fresh supplemented BP medium. At DIV11-16 neurons were transfected with indicated constructs using Lipofectamine 2000 (Invitrogen). Before transfection 300 µl conditioned medium was transferred to a new culture plate. For each well, 1.8 µg DNA was mixed with 3.3 µl Lipofectamine 2000 in 200 µl BP, incubated for 30 minutes at room temperature and added to the neurons. After 1 to 1.5 hours, neurons were briefly washed with BP and transferred to the new culture plate with conditioned medium with an additional 500 µl supplemented BP and kept at 37°C in 5% CO2 for 4-6 days.

### DNA constructs

GFP-CLCa was a gift from Dr. Blanpied. Halo-CLCa was obtained by replacing the GFP from GFP-CLCa for a Halo-tag using Gibson assembly (NEBbuilder HiFi DNA assembly cloning kit). GFP-CPG2 was obtained by replacing the HA-tag in the HA-CPG2 construct (gift from Dr. Nedivi) using Gibson assembly. GFP-Intersectin Long (Addgene plasmid # 47395) and GFP-CLCb (EED/QQN) (Addgene plasmid # 47422) were a gift from Peter McPherson. FCHO1-pmCherryC1 (Addgene plasmid # 27690), Epsin2-pmCherryC1 (Addgene plasmid # 27673), CALM-pmCherryN1 (Addgene plasmid # 27691), Amph1-pmCherryN1 (Addgene plasmid # 27692) and Syndapin2-pmCherryC1 (Addgene plasmid # 27681) were a gift from Christien Merrifield. FKBP-β2-adaptin-GFP (Wood et al., 2017) and HIP1R-GFP-FKBP (Addgene plasmid # 100752) were a gift from Stephen Royle. The AP2m1 patch 1 mutant (AP2m1-P1-HA) was a gift from Dr. Richmond (Raman et al., 2014). GFP-Syndapin I was a gift from Dr. Robinson. GFP-Eps15 was a gift from Dr. Van Bergen en Henegouwen The following constructs have been described before: Homer1c-mCherry, Homer1c-GFP, Dynamin2-GFP (Scheefhals et al., 2019), pSM155-mirItsn-GFP (van Gelder et al., 2020) , GFP-CLCa knock-in construct (Willems et al., 2020).

### Immunocytochemistry and HaloTag labelling

Neurons were fixed between DIV16-21 with 4% paraformaldehyde (PFA, EM grade) diluted in PEM buffer (80 mM PIPES, 5 mM EGTA, 2 mM MgCl2, pH 7.4) for 10 minutes at 37°C and washed three times with PBS supplemented with 100 mM glycine (PBS-gly). Then, neurons were permeabilized and blocked with 10% normal goat serum (NGS) and 0.01% Triton X-100 (TX) in PBS-gly for 30 minutes at 37°C. For STED imaging, GFP and mCherry containing constructs were enhanced with corresponding polyclonal anti-GFP (1:2000, MBL) and anti-mCherry (1:1000, Clonetech) antibodies diluted in PBS-gly supplemented with 5% NGS and 0.01% TX, for an overnight at 4°C. The next day, coverslips were washed three times in PBS-gly and anti-GFP was further labelled with ATTO647N (1:500, Sigma) and anti-mCherry was labelled with CF568 (1:500, Sigma) for 2 hours at room temperature (RT), washed and mounted in Mowiol (Sigma). For SMLM on Homer-mCherry and GFP-CLCa the same procedure was used as described above, but anti-GFP was labelled with Alexa-647-conjugated secondary antibodies (Life Technologies). After two hours, coverslips were washed three times and kept in PBS until further use. For SMLM on Halo-CLCa combined with various endocytic proteins fused to GFP, we first performed live-labeling with Halo-JF646 (1:1000, Promega) for 15 minutes at RT. To label endocytic proteins, GFP was labelled with a monoclonal anti-GFP (1:1000, Thermo Fisher) and labelled with a corresponding CF568-conjugated secondary antibody (Sigma). Although the localization density obtained for Halo-CLCa labelled with JF646 was lower compared to GFP-CLCa labelled with primary and secondary antibodies, no difference in CLCa morphology was observed (Supplement Figure 2).

### Confocal imaging

Confocal images were acquired with a Zeiss LSM 700 confocal laser-scanning microscope using a Plan-Apochromat 63x NA 1.40 oil objective. Images consist of a z-stack of 5-9 planes at 0.37-µm interval, and maximum intensity projections were generated in Fiji (Schindelin et al., 2012) for analysis and display.

### STED imaging

Gated STED (gSTED) images were taken with the Leica TCS SP83x microscope using a HC PL APO 100x/NA 1.4 oil immersion STED WHITE objective. The 488 nm pulsed white laser (80 MHz) was used to excite Alexa-488, 561 nm to excite CF568, and the 647 nm to excite JF646 and ATTO647N labeled proteins. JF646 and ATTO647N were depleted with the 775-nm pulsed depletion laser, and for depleting CF568 the 660-nm pulsed depletion laser was used. The internal Leica HyD hybrid detector was set at time gate between 0.3 and 6 ns. Images were taken with a pixel size lower than 40 nm, and Z-stacks were acquired. Maximum intensity projections were generated in Fiji (Schindelin et al., 2012) for analysis and display.

### Live-cell imaging

Live-cell imaging was performed on a spinning disk confocal system (CSU-X1-A1; Yokogawa) mounted on a Nikon Eclipse Ti microscope (Nikon) with Plan Apo VC 100x 1.40 NA with excitation from Cobolt Calyspso (491 nm), and Jive (561 nm) lasers, and emission filters (Chroma). The microscope was equipped with a motorized XYZ stage (ASI; MS-2000), Perfect Focus System (Nikon), Evolve 512 EM-CCD camera (Photometrics), and was controlled by MetaMorph 7.7.6 software (Molecular Devices). Neurons were maintained in a closed incubation chamber (Tokai hit: INUBG2E-ZILCS) at 37 °C in extracellular imaging buffer. For high frequency live-cell imaging (Figure 2A-D) images of GFP-CLCa were taken every 5 seconds for 5 minutes. For long-term live-cell imaging of Homer1c and CLCa (Figure 2E), images were taken every 30 seconds for 20 minutes. Lastly, imaging Homer1c and endocytic proteins fused to either mCherry or GFP was done taking images every 20 seconds for 10 minutes. In all the above-mentioned experiments Z-stacks of 5-9 planes were acquired, with varying step sizes per neuron. Also Homer1c-mCherry was only imaged in the first and last frame. Maximum intensity images were analyzed in Fiji, by manually drawing same-size ROIs around individual puncta associated with PSDs. To measure lifetimes of clathrin and endocytic proteins we used the TrackMate plugin (Tinevez et al., 2017).

### Fluorescence recovery after photobleaching

FRAP experiments were performed on the spinning disk confocal system as described above, using the ILas2 system (Roche scientific). A baseline of 2 minutes with a 20-second interval was taken, followed by photobleaching of individual puncta with a targeted laser. The recovery of fluorescence of GFP-CLCa was imaged for 3 minutes with 20-second interval, followed by 12 minutes with 60-second interval, and 4 minutes with 120-second interval, resulting in a total recovery time of 19 minutes. For imaging the recovery of the endocytic proteins an acquisition of 10 minutes was taken (2-minute baseline, 3 minutes with 20-second interval and 7 minutes with 60-second interval). For acquiring FRAP images, a single Z-plane was taken. Fluorescence intensity was measured in Fiji, by manually drawing same-size ROIs around puncta. For analysis, acquisitions were corrected for drift. For each ROI, the mean intensity was measured for every time point and corrected for background and bleaching. Normalized intensities were plotted over time. Individual curves were fitted with a single-exponential function I = A(1 – exp^(-Kt)^) to estimate the mobile fraction (A) and time constant tau.

### Single-molecule localization microscopy and analysis

dSTORM data was acquired on the Nanoimager S from ONI (Oxford Nanoimaging Ltd.), equipped with a 100x, 1.4NA oil immersion objective, an XYZ closed-loop piezo stage, and four laser lines: 405-nm, 471-nm, 561-nm and 640-nm. Fluorescence emission was detected using a sCMOS camera (ORCA Flash 4, Hamamatsu). Stacks of 10,000 images were acquired at 20 Hz in TIRF mode. Samples were imaged in PBS containing 10 - 50 mM MEA, 5% w/v glucose, 700 μg/ml glucose oxidase, and 40 μg/ml catalase. Data was processed in NimOS software from ONI. Before each imaging session, a bead sample calibration was performed to align the two channels, achieving a channel mapping precision smaller than 8 nm. Images were rendered in ONI software and loaded into Fiji. Here, ROIs of 1 x 1 µm were drawn around individual EZ. The ROI sets were imported in Matlab (2018b) for analysis.

First, tracking was performed on the localization data to merge localizations that were detected in more than two consecutive frames as described in (Willems et al., 2020) . Next, a localization cutoff of 15 nm was taken to further analyze the localization data. A DBScan was performed to define the borders of Homer1c and CLCa in figure 4, using an epsilon of 0.2 and minimum number of localizations of 100. For figure 6, an epsilon of 0.35 and minimum number of localizations of 50 was used. For figure 6, rings were applied to reveal the relative distribution of endocytic proteins to the EZ. Rings were calculated as a percentage of the 100% polyshape given by the DBScan. Inwards, 5 rings were created: 0-20, 20-40, 40-60, 60-80, 80-100; and outwards 3 rings were created: 100-120, 120-140, 140-160. Then, the number of localizations for each of the endocytic proteins were calculated per ring. The fraction of these localizations per EZ, were plotted against the fraction of the area of the ring and normalized to 1.

### Quantification of EZ-associated synapses

For the Shank knockdown experiments DIV14 neurons were transfected with pSM155-mCherry or pSM155-mirShank-mCherry together with Homer1c-ALFA and indicated construct. Homer1c-ALFA was labelled with JF646-conjugated FluoTAG X4 anti-ALFA (1:500 Fluotag X4, Nanotag). In the experiments manipulating actin dynamics, Homer-mCherry and GFP-CLCa expressing neurons were incubated with Latrunculin B (20 µM, Bioconnect), CK666 (400 µM, Tocris) or Jasplakinolide (20 µM, Tocris) in E4 for 30 minutes at 37°C and fixed immediately after. As a control E4 containing DMSO was used. GFP-CLCa or GFP-CLCb-EED/QQN were co-expressed with Homer1c-mCherry. pSM155-mirItsn was co-expressed with Homer1c-mCherry and Halo-CLCa labelled with JF646. AP2m1-WT or AP2m1-P1-HA was co-expressed with Homer1c-mCherry and GFP-CLCa. To quantify the fraction of synapses with an associated EZ or puncta of endocytic protein, circular regions with a fixed diameter (0.69-0.89 µm) were centered on the Homer1c signal to outline synaptic regions. These regions were then transferred to the GFP-CLC or tagged endocytic protein channel. A synapse was classified positive if the endocytic protein cluster overlapped partially or completely with the circular region. The fraction of positive synapses was calculated per cell and averaged per condition over the total population of neurons. Data plotted is normalized to the average of the control.

### Statistical analysis

Statistical significance was tested using a student’s t-test when comparing two groups. When comparing multiple groups statistical significance was tested using a one-way ANOVA followed by a Tukey or Dunnett’s multiple comparison post-hoc test. All the statistical tests with a p-value below 0.05 were considered significant. In all figures, significance is indicated as follows: p < 0.05 is indicated by *, p < 0.01 by **, and p < 0.001 by ***. Analysis was performed on neurons originating from at least two individual batches of hippocampal neurons. Number of neurons used for analysis is indicated as *N*, number of spines or clathrin-coated structures is represented as *n*.

## ACKNOWLEGDEMENTS

We would like to thank all members of the MacGillavry lab for support and discussions. This work was supported by the Netherlands Organization of Scientific Research (NWO-ALWOP. 191 to H.D.M).

## AUTHOR CONTRIBUTION

Conceptualization, Methodology, Validation & Format Analysis: L.A.E.C and H.D.M; Investigation: L.A.E.C., M.W., A.M.L.S; Resources: H.D.M; Writing and Editing: L.A.E.C and H.D.M; Supervision: H.D.M; Funding: H.D.M

---

**Supplementary Figure 1:**
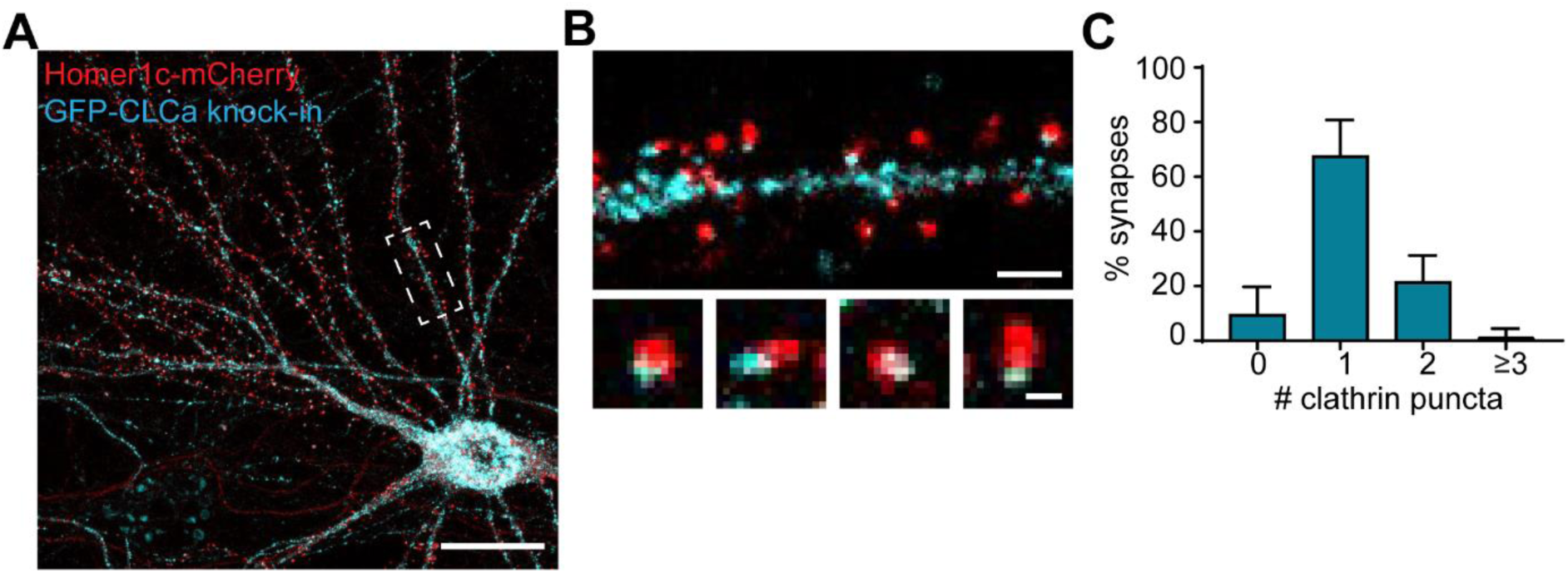
Distribution of endogenously tagged CLCa in neurons. (A) Example image of neuron expressing Homer1c-mCherry and GFP-CLCa knock-in construct. Scale bar: 20 µm. (B) Example image of clathrin-coated structures in the dendrite (upper panel) and zooms of individual synapses associated with an EZ (lower panel). Scale bars: 2 µm, zooms: 500 nm. (C) Quantification of the average number of clathrin-coated structures per synapse, plotted as mean ± SEM (n = 6).

**Supplementary Figure 2.**
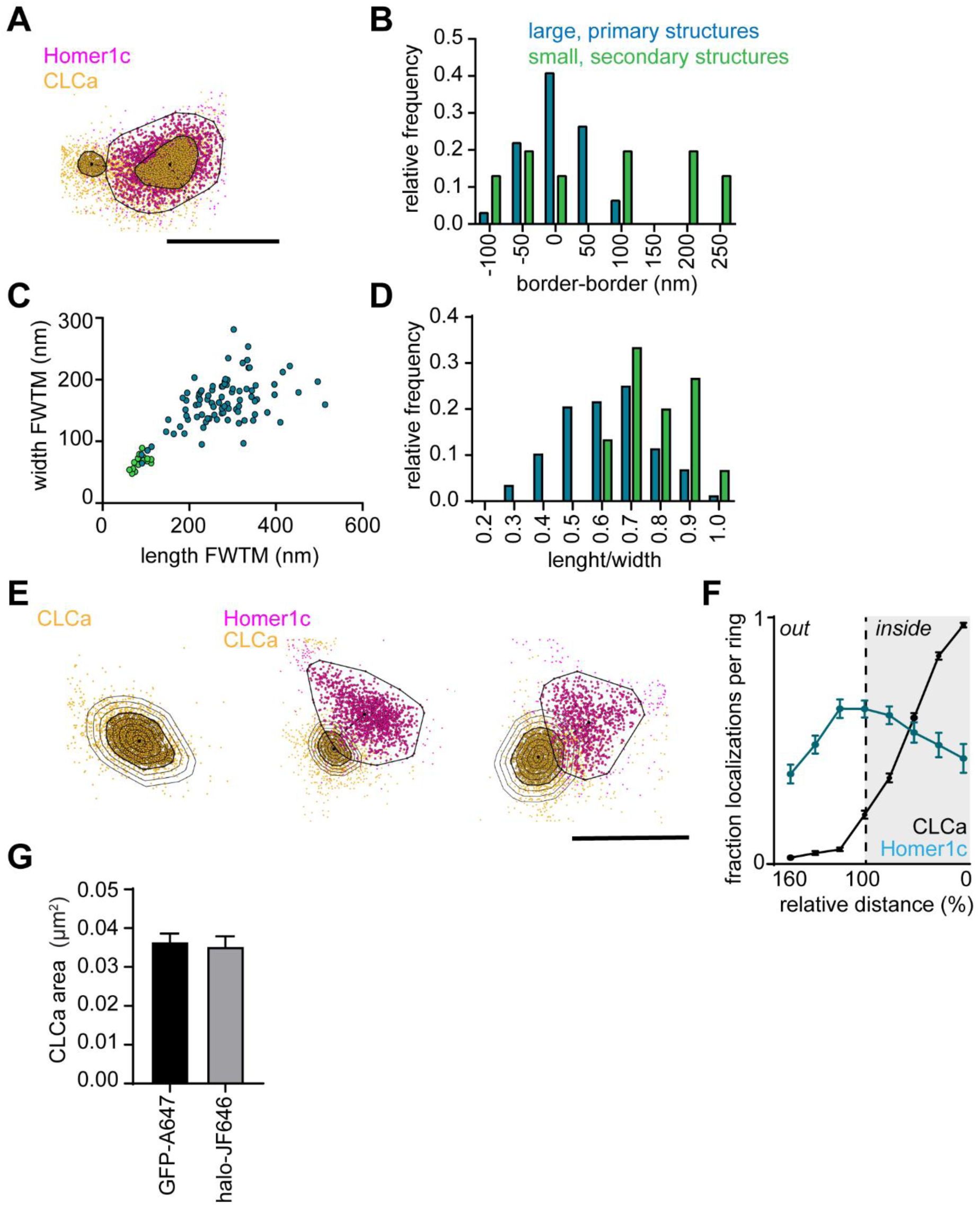
SMLM reveals nanoscale scale architecture of PSD-associated CLCa structures. (A) Example plots visualizing the individual localizations obtained for Homer1c (magenta) and CLCa (orange). Two or more individual CLCa structures can be observed per PSD. Scale bar: 500 nm. (B-D) Data visualized in these plots is the same data represented in Figure 3, but here a distinction was made between PSDs containing one (blue) or two (green) CLCa structures is made. (B) Histogram of the distance from the border of the PSD to the border of CLCa structures. (C) Scatterplot showing the dimensions of larger (blue) CLCa structures, most likely representing the EZ, and smaller CLCa structures (green) (big structures: n = 92, small structures: n = 13). (D) Histogram showing circularity ratio of CLCa structures. (E) Example plots of CLCa with rings based on the size of the structure (left panel), and example plots of individual CLCa and Homer1c localization. Scale bar: 500 nm. (F) Fraction of localizations per ring. Dotted line depicts the border of CLCa (Homer1c n = 65, CLCa n = 66). (G) Comparison of the average area of CLCa puncta labelled with different strategies (GFP-CLCa labelled with monoclonal GFP and CF568, Halo-CLCa labelled with halo-JF646), plotted as mean ± SEM (GFP-CF568: n = 103, Halo-JF646: n = 50).

**Supplementary Figure 3.**
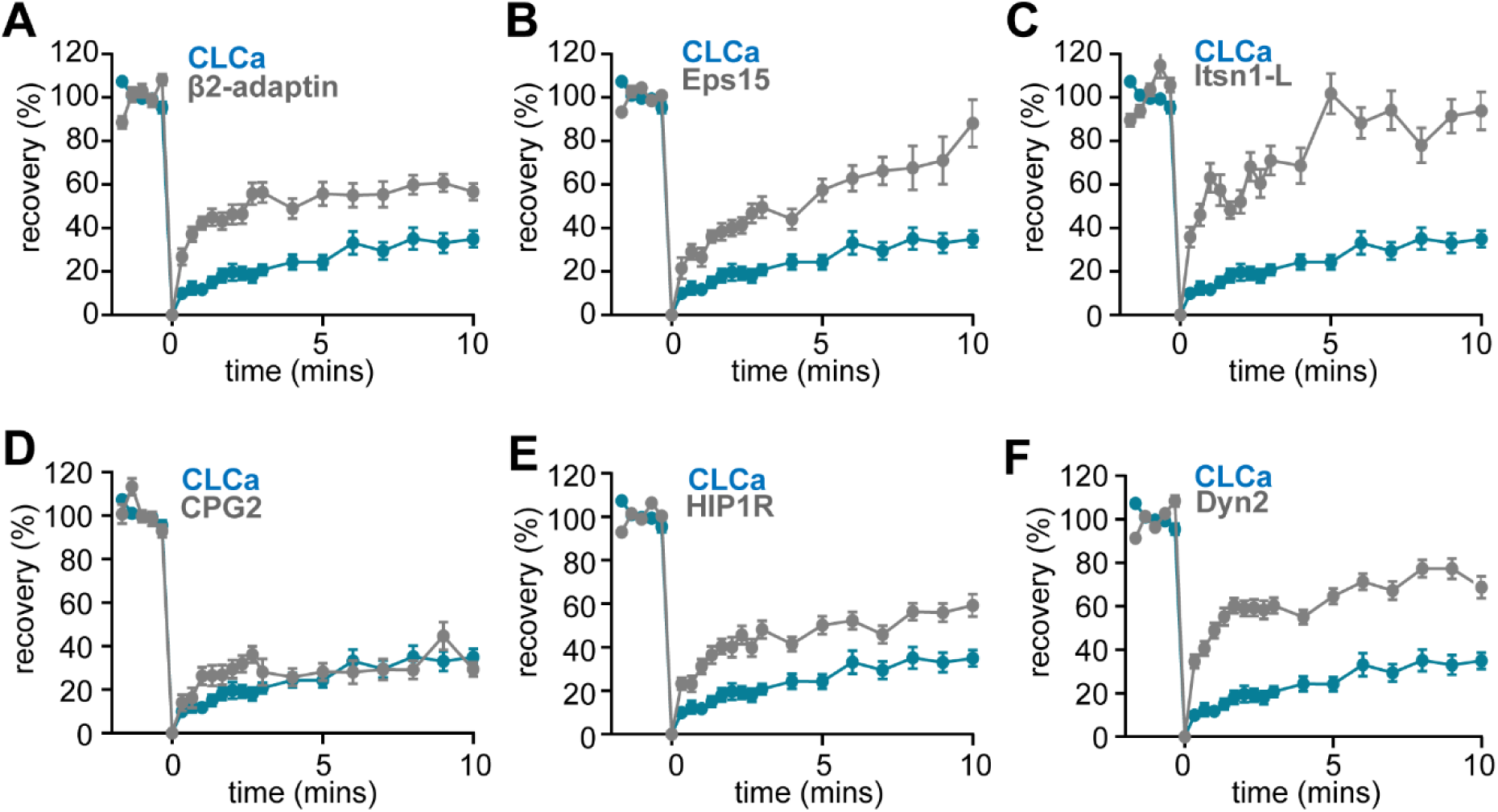
FRAP curves of endocytic proteins compared to CLCa. (A-G) FRAP curves of endocytic proteins (grey) and CLCa (blue) after bleaching on t = 0 for (A) β2-adaptin (n = 13), (B) Eps15 (n = 23), (C) Itsn1L (n = 20), (D) CPG2 (n = 22), (E) HIP1R (n = 44) and (F) Dyn2 (n = 51). All graphs show the same CLCa (n = 32) data for comparison. All data is plotted as mean ± SEM.

**Supplementary Figure 4.**
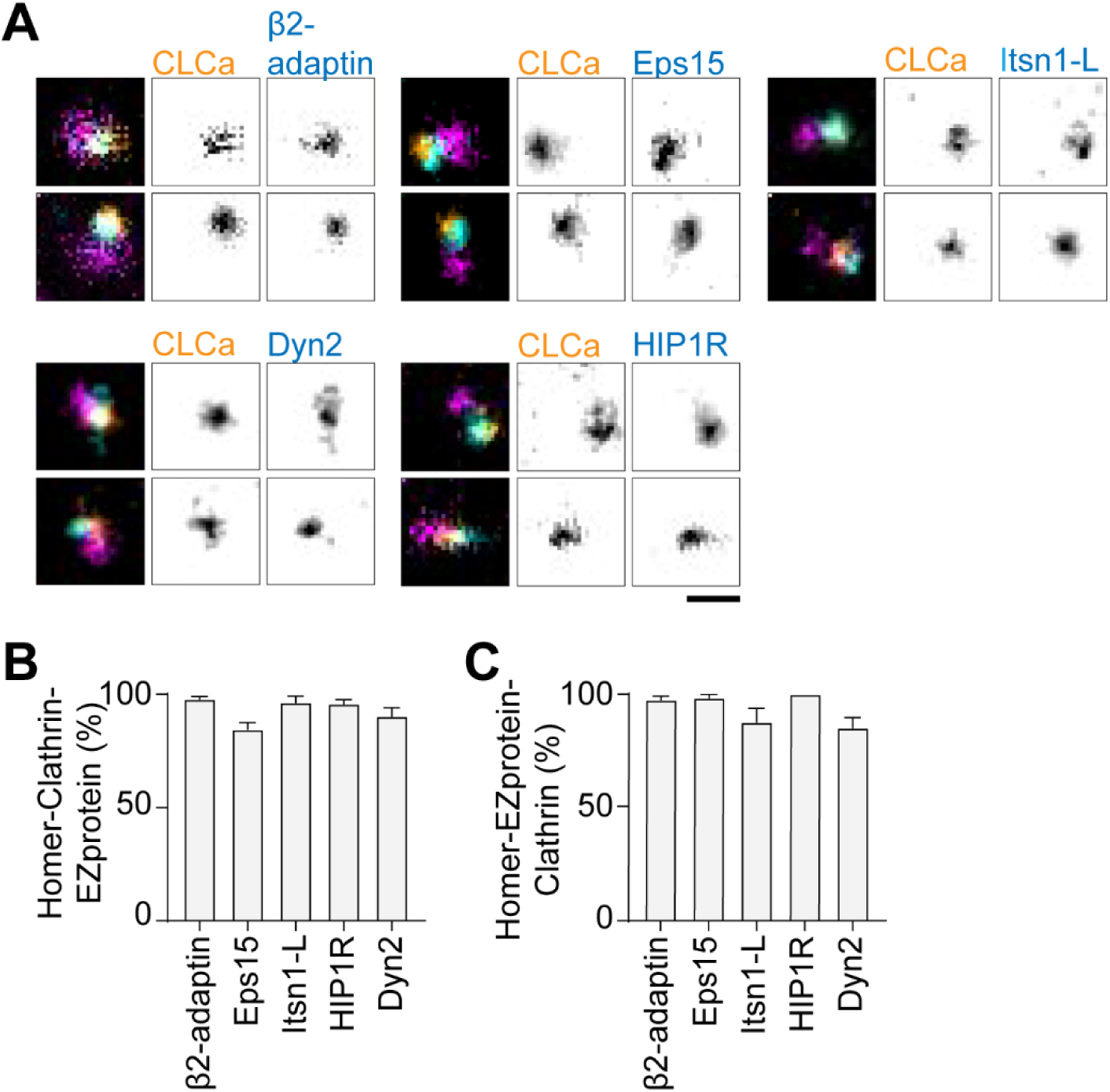
Endocytic proteins colocalize with the EZ. (A) Examples of endogenous Homer1b/c (eHomer1b/c) labelled with anti-Homer1 antibody (magenta), combined with Halo-CLCa labelled with JF646 (orange) and co-expressed with endocytic proteins fused to GFP labelled with CF568 (cyan). eHomer1b/c was imaged using confocal, for CLCa and endocytic proteins gSTED was applied. Scale bar: 500 nm. (B) Percentage of eHomer1c-associated CLCa puncta overlapping with endocytic protein signal. (C) Percentage of eHomer1c-associated endocytic protein puncta overlapping with CLCa signal. Bar graphs show mean ± SEM, normalized to the average of the representative controls described in method section.

